# Mass spectrometry-based top-down proteomics in nanomedicine: proteoform-specific measurement of protein corona

**DOI:** 10.1101/2024.03.22.586273

**Authors:** Seyed Amirhossein Sadeghi, Ali Akbar Ashkarran, Morteza Mahmoudi, Liangliang Sun

**Author notes:** Corresponding Authors. Morteza Mahmoudi Liangliang Sun.

## Abstract

Conventional mass spectrometry (MS)-based bottom-up proteomics (BUP) analysis of protein corona [i.e., an evolving layer of biomolecules, mostly proteins, formed on the surface of nanoparticles (NPs) during their interactions with biomolecular fluids] enabled nanomedicine community to partly identify the biological identity of NPs. Such an approach, however, fails pinpoint the specific proteoforms—distinct molecular variants of proteins, which is essential for prediction of the biological fate and pharmacokinetics of nanomedicines. Recognizing this limitation, this study pioneers a robust and reproducible MS-based top-down proteomics (TDP) technique for precisely characterizing proteoforms in the protein corona. Our TDP approach has successfully identified hundreds of proteoforms in the protein corona of polystyrene NPs, ranging from 3-70 kDa, revealing over 20 protein biomarkers with combinations of post-translational modifications, signal peptide cleavages, and/or truncations—details that BUP could not fully discern. This advancement in MS-based TDP offers a more comprehensive and exact characterization of NP protein coronas, deepening our understanding of NPs’ biological identities and potentially revolutionizing the field of nanomedicine.

## Introduction

Nanomedicine applies nanotechnology concept to medicine, i.e., employing biocompatible nanoparticles (NPs) for controlled and/or targeted delivery of therapeutic (bio)molecules to desired tissues/organs, imaging, and disease diagnosis.^1–8^ The overall efficacy of nanomedicine is strongly impacted by protein/biomolecular corona, i.e., the composition and decoration of various types of biomolecules (e.g., mostly proteins) that bound to the surface of NPs after they are exposed to biological fluids (e.g., human plasma).^9,10^ It has been well documented that the composition and decoration of participated proteins in protein corona determines the biological fate and pharmacokinetics of NPs.^11–16^ Therefore, achieving comprehensive and accurate characterization of the composition of protein corona is central to advance nanomedicines’ safety together with their therapeutic and diagnostic efficacy.^9^ In addition, protein corona has also been recognized as a useful analytical technique to discover new protein biomarkers of diseases because it can reduce the complexity of biological fluids (e.g., plasma), which enables easier detection and identification of biomarkers.^17–23^

Mass spectrometry (MS)-based bottom-up proteomics (BUP) has been widely recognized as an efficient way for the characterization of protein corona, providing the identification of gene products in the protein corona.^9,19,24^ However, MS-based BUP fails to identify exact forms of protein molecules (i.e., proteoforms^25^) in the protein corona because of the “peptide-to-protein” inference problem.^26^ Proteoforms from the same gene could have divergent biological functions^27–30^ and proteoforms are vital for modulating disease progression^31–35^. Therefore, proteoform-specific measurement of protein corona will undoubtedly provide more accurate characterization of protein molecules in the protein corona, not only bettering our understanding of how protein corona directs the interactions NPs in biological systems (e.g., immune cells) but also offering new opportunities for novel proteoform biomarker discovery.

MS-based top-down proteomics (TDP) directly measures intact proteoforms in complex biological samples via coupling high-resolution proteoform separations [i.e., liquid chromatography (LC) and capillary electrophoresis (CE)] with advanced MS and tandem MS (MS/MS).^36^ It is an ideal strategy for the characterization of proteoforms and provides a bird’s-eye view of proteoforms in cells, tissues, and biofluids under various biological conditions.^27,29–31,37^ Here, for the first time, an efficient and reproducible TDP technique was developed for the proteoform-specific measurement of protein corona.

## Results

### A MS-based TDP workflow for the characterization of protein corona

To develop an efficient TDP workflow for protein corona, we employed polystyrene NPs (PSNPs) and a commercially available healthy human plasma sample to form protein corona and utilized dynamic pH junction-based capillary zone electrophoresis (CZE)-MS/MS^38–40^ for online proteoform separation, detection, and identification. **Figure 1** illustrates the detailed TDP workflow. PSNPs were incubated with a human plasma sample to form protein corona. The protein corona was then eluted from the PSNP surface using an elution buffer containing 0.4% (w/v) SDS. The eluted protein corona sample was buffer-exchanged to a MS-compatible buffer (100 mM ammonium bicarbonate, pH 8),^41^ followed by dynamic pH junction-based CZE-MS/MS analysis using an Orbitrap Exploris 480 mass spectrometer.

**Figure 1.**
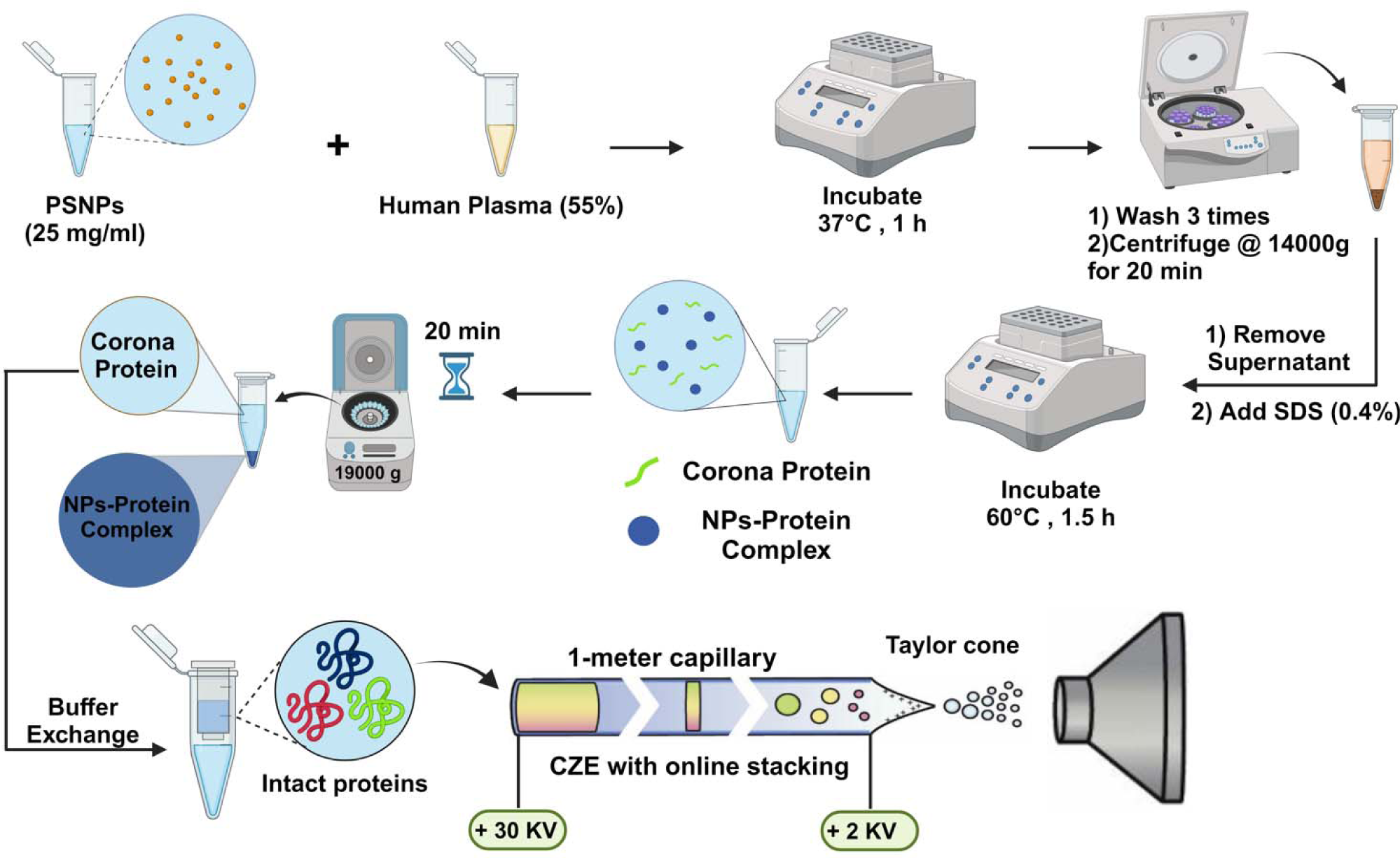
Schematic of the MS-based TDP workflow for protein corona using polystyrene nanoparticles (PSNPs), a human plasma sample, and capillary zone electrophoresis (CZE)-tandem MS (MS/MS).

Both “high-high” and “low-high” MS modes^42^ were used to measure proteoforms in the protein corona. In the “high-high” mode, proteoform parent ions (MS) were detected using a 480,000 resolution (at m/z 200) and fragment ions (MS/MS) were measured using a 120,000 resolution (at m/z 200). The “high-high” mode was mainly used to identify proteoforms smaller than 30 kDa. In the “low-high” mode, proteoform parent ions (MS) were detected using a 7,500 resolution (at m/z 200) and fragment ions (MS/MS) were still measured using a 120,000 resolution (at m/z 200). Since the 480,000 resolution (at m/z 200) still has difficulties in achieving isotopically resolved peaks for proteoforms larger than 30 kDa, the low-resolution MS was employed to measure the large proteoforms for determining their average masses. Finally, the TopPIC software developed by Liu’s group^43^ was used for database search to identify and quantify proteoforms from the “high-high” mode. For the “low-high” mode data, we employed the UniDec software^44^ for mass deconvolution to determine the average masses of large proteoforms and employed the ProSight Lite bioinformatics tool^45^ to perform the identification of large proteoforms based on the masses of proteoforms and corresponding fragments in a targeted fashion.

### Efficient and reproducible TDP measurements of protein corona

Protein corona was formed on the surface of PSNPs after incubating the PSNPs and human plasma sample. The particle size increased from 78.9 ± 0 nm to 105.3 ± 3.8 nm after the formation of protein corona based on the dynamic light scattering (DLS) measurement. The particle size distribution is reasonably narrow according to the standard deviation (SD) and polydispersity index (PDI) data (see **Figure S1A** for more details). Interestingly, the particle-size SD and PDI of PSNPs are higher after protein binding compared to the bare NPs (**Figure S1A** and **S1B**), suggesting some extent of heterogeneity of protein corona on individual PSNPs. After protein corona elution, the size of particles decreased to 92.9 ± 0 nm and the PDI of 0.042, suggesting that the elution procedure using a 0.4% (w/v) SDS solution can efficiently elute proteins in the outer layer of protein corona. The zeta potential of PSNPs has a substantial change after protein binding and protein elution compared to bare NPs, agreeing with the particle size data, **Figure S1A**. Transmission electron microscopy (TEM) was further employed to characterize the PSNPs before protein binding (bare NPs), after protein binding, and after protein elution, **Figure 2A**. After incubating with human plasma, the surface of PSNPs was clearly covered by a darker shell, indicating the formation of protein corona. After protein elution, there are much less proteins on the PSNPs. All the measurement data on PSNPs demonstrate that protein corona was successfully formed on the surface of PSNPs and the protein elution procedure is sufficient for eluting the outer layer of protein corona.

**Figure 2.**
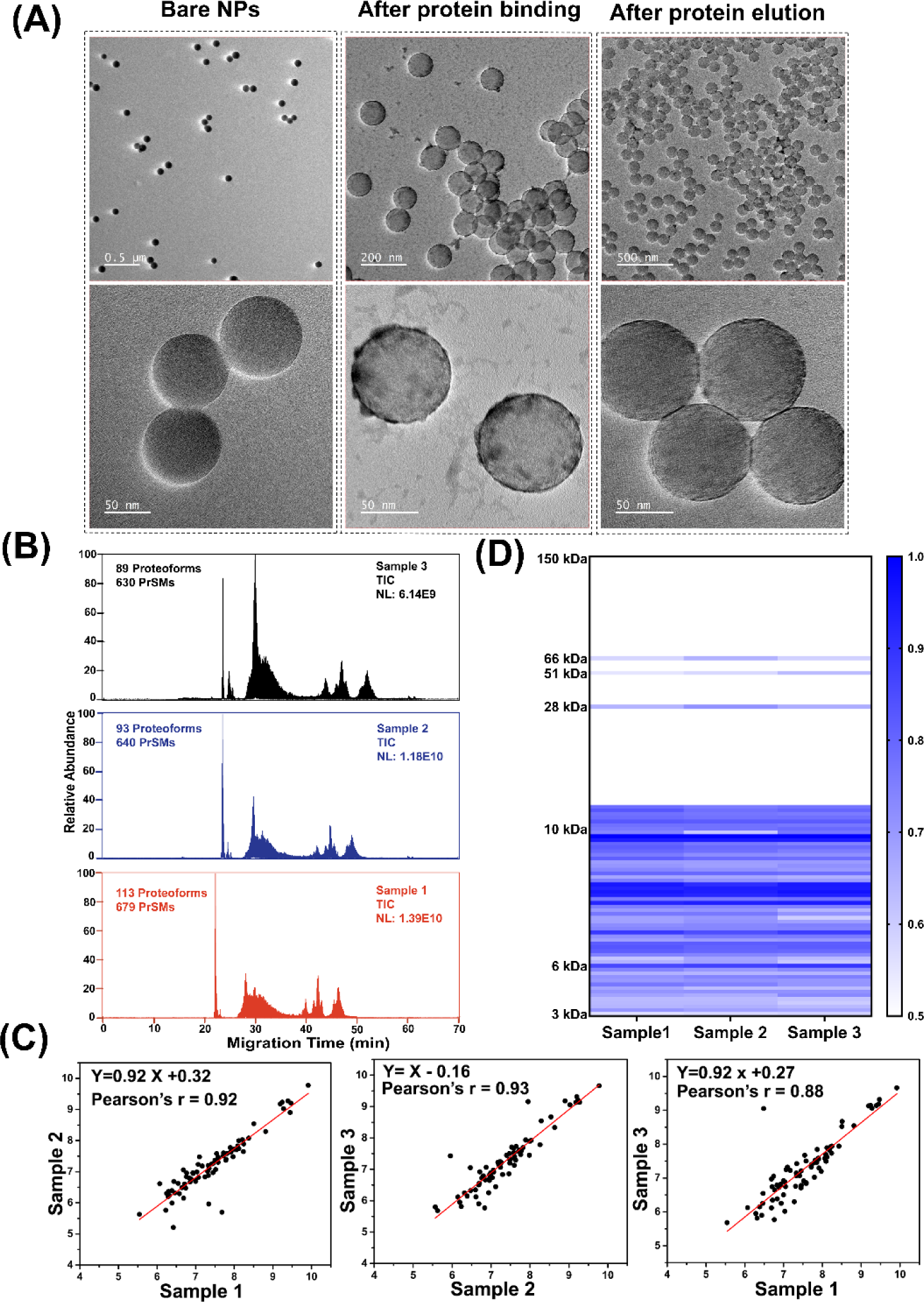
Characterization of protein corona. (A) TEM images of Bare NPs, NPs after protein binding, and NPs after protein elution with 0.4% SDS. (B) Examples of total ion current (TIC) electropherograms of eluted protein corona after CZE-MS/MS analyses in “high-high” mode. Three protein corona samples (sample 1-3) were prepared in parallel and analyzed by CZE-MS/MS. Each sample was measured in triplicate. (C) Proteoform intensity correlations between any two samples. The data are from “high-high” mode. Log-log plots are shown in the figure. (D) Heatmap of detected proteoform intensity across three samples. Proteoform intensity was log2 transformed and used to create the heat map using GraphPad Prism.

The eluted protein corona sample was measured by SDS-PAGE (**Figure S2A**) and CZE-MS/MS (**Figure 2B-D** and **Figure S2B**). Three samples were prepared in parallel starting from the PSNP and human plasma incubation for evaluating the overall reproducibility of the TDP technique. The SDS-PAGE data shows consistent proteoform profiles across the three samples with strong bands at around 25 kDa and between 50-75 kDa. The CZE-MS/MS data (“high-high” mode) also show consistent total ion current (TIC) electropherograms, the number of proteoform identifications (98±13), and the number of proteoform-spectrum matches (PrSMs, 650±26) across the three samples, **Figure 2B**.

To assess the quantitative reproducibility of CZE-MS/MS, we examined the correlation coefficients of proteoform intensities between any two samples, **Figure 2C**. The proteoform intensity was obtained using the TopDiff software^46^ from the “high-high” mode data. The shared proteoforms among any two samples were utilized for the analysis. The intensities of the shared proteoforms from technical triplicate measurements were averaged and used to generate the plots. The strong linear correlations (Pearson’s *r* = 0.88-0.93) indicate the high quantitative reproducibility of the TDP technique developed in this work for measuring proteoforms in the protein corona samples. The proteoform intensity heatmap in **Figure 2D** further illustrates the reproducibility of our TDP technique in measuring proteoforms (3-70 kDa) across three protein corona samples prepared in parallel. The CZE-MS/MS in “high-high” mode only enabled the identification of proteoforms around 10 kDa or smaller. The proteoforms larger than 28 kDa in **Figure 2D** were from the “low-high” mode.

Figure S2B shows the consistent base peak electropherogram profiles of the three protein corona samples in the “low-high” mode. Subsequently, extracted ion electropherograms (EIEs) of three large proteins (∼28 kDa, ∼51 kDa, and ∼66 kDa) from the three samples were obtained, Figure 3A. The separation profiles and peak intensities of the three proteins are relatively consistent across the three samples. The peak 1 shows obvious intensity variations across the three samples, which may be due to the limited number of data points across the top part of the peak, arising from the relatively long data acquisition cycle in TDP measurement. Figure 3B shows the mass spectra and deconvoluted masses of the three large proteins. Interestingly, multiple proteoforms for each protein were detected and the deconvoluted data shows the masses and relative abundance of different proteoforms in each protein peak. For example, three clear proteoforms of protein 1 with masses 66,560 Da, 66,872 Da (66,560+312 Da), and 67,184 Da (66,560+312+312 Da) were observed with the 66,560-Da proteoform as the most abundant one. Based on the mass and the list of proteins in the corona sample from BUP measurement, we presumably identified the protein as human serum albumin (HSA). The MS/MS data also provides strong support about the identification of the HSA with a series of matched b-type fragment ions close to the N-terminus of the protein, **Figure S3**. The theoretical mass of HSA is 66,438 Da considering the 17 disulfide bonds (native form). The most abundant proteoform (66,560 Da) represents a +122-Da mass shift compared to the native form, presumably due to a combination of one phosphorylation (+80 Da) and one acetylation (+42 Da) or cysteinylation (+119 Da)^47^. The other two proteoforms of HSA most likely have additional glycosylation PTMs according to the literature data and the information in the UniProt knowledgebase (https://www.uniprot.org/uniprotkb/P02768/entry).

**Figure 3.**
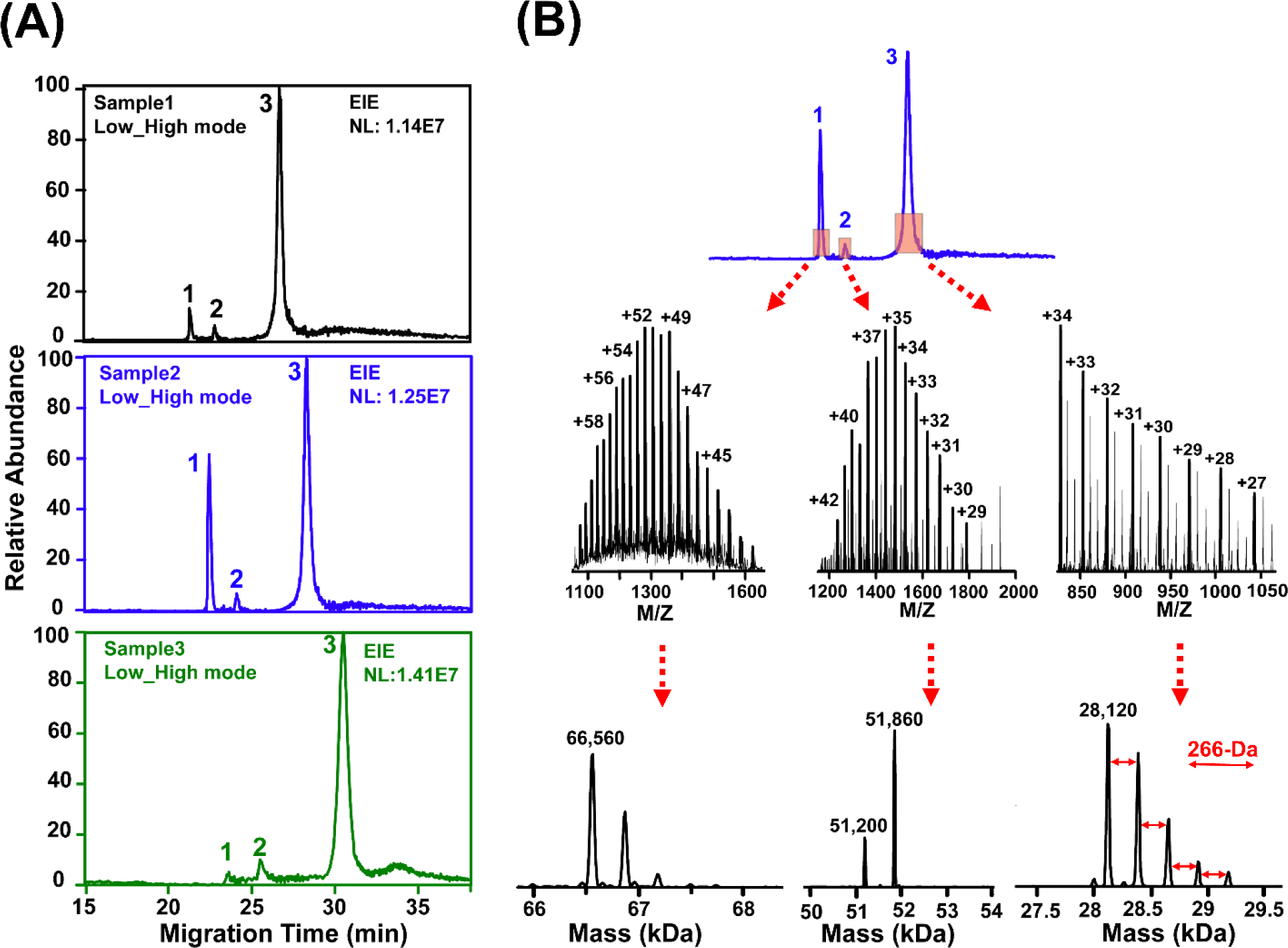
Measurement of large proteoforms. (A) Extracted ion electropherograms (EIE) of three large proteins (1-3). The m/z ion with the highest intensity for each protein was used for the peak extraction with a mass tolerance of 200 ppm. (B) Averaged mass spectrum of each protein across the peak and the corresponding deconvoluted masses of various proteoforms. The mass deconvolution was performed using the UniDec (Universal Deconvolution) software with default settings.^44^

For the 28-kDa protein in Figure 3B, five proteoforms were clearly detected and the most abundant one has a mass of 28,120 Da. According to the proteoform mass and our BUP measurement results, the protein was presumably identified as Apolipoprotein A-I (APOA1). The MS/MS data offers strong evidence about the identification with a series of matched y-type fragment ions close to the C-terminus of the protein, **Figure S4**. Interestingly, the five different proteoforms of intact APOA1 show a continuous increase in proteoform mass from the highest abundant to the lowest abundant one with a 266-Da mass difference between any neighboring proteoforms. The 266-Da mass difference could be due to lipidation, for example, adding a stearic acid (octadecanoic acid, 284 Da) molecule through reaction between the carboxyl group of lipid and amine group of protein via removing a H_2_O molecule. The five different proteoforms could represent APOA1 with 0-4 stearic acid modification sites. One previous TDP study provided strong evidence that APOA1 proteoforms are closely related to the indices of cardiometabolic health.^48^ APOA1 is a prognostic marker in renal and liver cancers according to the human protein atlas (https://www.proteinatlas.org/ENSG00000118137-APOA1). The data here suggests that coupling PSNP-based protein corona and MS-based TDP could be a valuable strategy for discovering novel APOA1 proteoform biomarkers of diseases (i.e., cancers and cardiometabolic diseases) using human plasma samples.

For the 51-kDa protein, two proteoforms were detected with 51,200 Da and 51,860 Da. Based on the mass and our BUP data of the protein corona sample, the protein might be Fibrinogen beta chain (50,763 Da in the mature form without PTMs) or Clusterin (50,062 Da in the mature form without PTMs). Unfortunately, the MS/MS spectra of those two detected proteoforms do not match well with theoretical b-or y-type of fragment ions from Fibrinogen beta chain or Clusterin. More additional studies are needed in the future to provide more information about the identities of those two proteoforms.

### Comparisons between TDP and BUP for the characterization of protein corona

The protein corona samples were analyzed by both BUP and TDP using CZE-MS/MS. For the BUP, the protein corona on PSNPs was prepared through the on-bead digestion procedure in the literature.^19,24^ For TDP, the protein corona was eluted from PSNPs and cleaned prior to CZE-MS/MS analysis, as shown in Figure 1. Three protein corona samples prepared in parallel starting from the same human plasma sample were analyzed.

Figure 4 summarizes the protein corona data from TDP (Figure 4A) and BUP (Figure 4B). Single-shot TDP analysis of the protein corona sample by CZE-MS/MS consistently identified nearly 600 PrSMs, 100 proteoforms, and 20 proteoform families across the three protein corona samples, Figure 4A. The Proteoform family represents a set of proteoforms from the same gene.^49^ In total, 263 proteoforms and 50 proteoform families were identified from the three corona samples. Single-shot BUP analysis of the protein corona sample by CZE-MS/MS identified around 8,000 peptide-spectrum matches (PSMs), 3,000 peptides, and 200 protein groups with high reproducibility, Figure 4B. A protein group can contain multiple proteins sharing the same set of identified peptides and those proteins usually have similar protein sequences. BUP analyses of the three protein corona samples using CZE-MS/MS identified, in total, 280 protein groups and 5,606 peptides.

**Figure 4.**
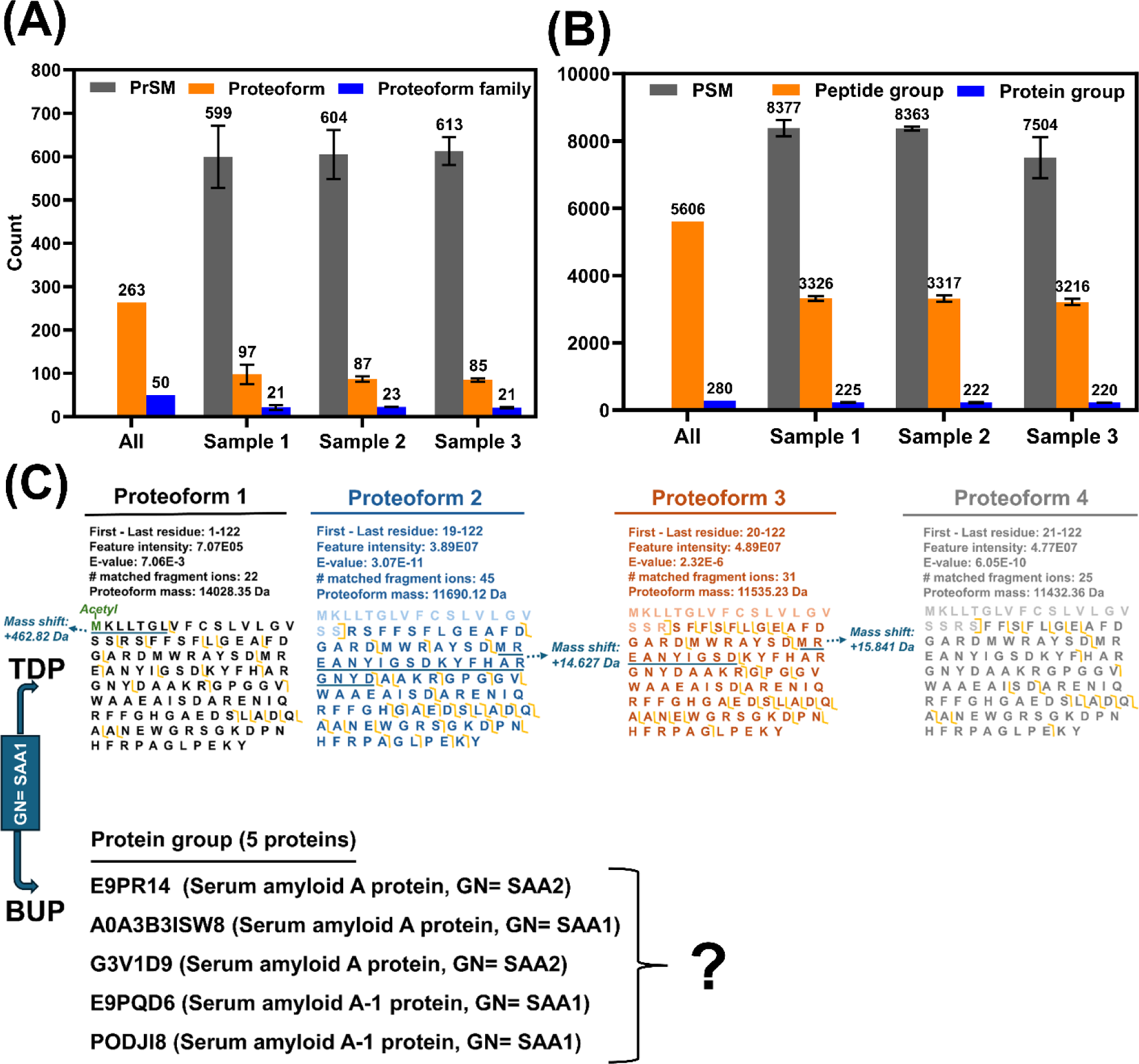
Differences between TDP and BUP for the characterization of protein corona. (A) The numbers of proteoform identifications, proteoform-spectrum matches (PrSMs), and proteoform family identifications from TDP. (B) The numbers of peptide identifications, peptide-spectrum matches (PSMs) and protein group identifications from BUP. (C) Four example proteoforms of *SAA1* identified by TDP with sequences and fragmentation patterns, and the five proteins included in the protein group SAA1 identified by BUP.

TDP produced a much lower proteome coverage than BUP in terms of the number of identified genes (50 *vs.* ∼280) due to its drastically lower sensitivity, resulting from much wider charge state distributions of intact proteoforms compared to peptides after electrospray ionization.^50^ However, TDP offered more precise and more valuable measurements of proteins in a proteoform-specific manner. For example, 18 different proteoforms of the gene Serum amyloid A-1 (*SAA1*) were identified by CZE-MS/MS-based TDP (“high-high” mode) from the protein corona samples. The SAA1 protein is a prognostic marker of renal cancer according to the Human Protein Atlas (https://www.proteinatlas.org/ENSG00000173432-SAA1). Four of the identified *SAA1* proteoforms are shown in Figure 4C. Those proteoforms were identified with high confidence evidenced by the E-Value and the number of matched fragment ions. They have varied length of sequences due to variations in signal peptide cleavage. For example, proteoform 1 has the whole protein sequence without signal peptide cleavage; proteoforms 2-4 have signal peptide cleavage at slightly different positions. Additionally, various PTMs occurred on those proteoforms. For example, proteoform 1 carries a N-terminal acetylation and one +463-Da mass shift close to the N-terminus; proteoform 2 has a roughly 14-Da mass shift, which could be a methylation PTM; proteoform 3 contains a 16-Da mass shift, most likely corresponding to an oxidation modification; proteoform 4 does not have any PTMs. The location of those PTMs on the proteoforms is at the underlined regions. The TDP approach also enabled the determination of relative abundance of proteoforms from the same gene. For example, proteoform 1 has a much lower abundance than the other three proteoforms, suggested by their intensities. On the other hand, the BUP measurement identified a protein group SAA1, containing five proteins, which have similar protein sequences. The main issue of the BUP data is that the protein identification is ambiguous, which means which protein (s) in the protein group exist in the protein corona sample is unresolved due to the “peptide-to-protein” inference problem.^26^

Additional data analyses were performed in terms of the identified protein biomarkers in this study. TDP identified specific proteoforms of 23 protein biomarkers, **Table 1**. The number of proteoforms ranges from 1 to 51 for those protein biomarkers with an average of 10 proteoforms per protein biomarker. The BUP data of those protein biomarkers show that each protein group can have a range of 1-22 proteins with an average of nearly 7 proteins per protein group. The data further indicates the advantages of TDP for protein analysis with more precise and more valuable proteoform measurements compared to BUP.

**Table 1.** Summary of some disease-related protein biomarkers identified by TDP and BUP.^#^

| Gene | TDP | BUP |  |
| --- | --- | --- | --- |
|  | # of Proteoform | Protein Group | # of Protein |
| APOC3 | 28 | Apolipoprotein C-III | 3 |
| APOC2 | 38 | Apolipoprotein C-II | 5 |
| APOC1 | 11 | Apolipoprotein C-I | 6 |
| APOB | 2 | Apolipoprotein B-100 | 7 |
| APOA2 | 51 | Apolipoprotein A-II | 6 |
| APOA1 | 49 | Apolipoprotein A-I | 2 |
| ACTN1 | 1 | Actinin alpha 1 | 22 |
| *TLN2 | 1 | Talin-1/2 | 4 |
| SERPINA1 | 11 | Alpha-1-antitrypsin | 15 |
| TTR | 1 | Transthyretin | 2 |
| ITGB3 | 1 | Integrin beta 3 | 3 |
| *QSOX1 | 2 | Sulfhydryl oxidase 1 | 2 |
| CLU | 1 | Clusterin | 10 |
| DCTN1 | 1 | Dynactin subunit 1 | 5 |
| ALB | 1 | Albumin | 8 |
| FGB | 1 | Fibrinogen beta chain | 2 |
| FGA | 2 | Fibrinogen alpha chain | 1 |
| SAA1 | 18 | Serum amyloid A-1 | 5 |
| C3 | 14 | Complement C3 | 10 |
| SHBG | 2 | Sex hormone-binding globulin | 13 |
| SERPINA6 | 1 | Corticosteroid-binding globulin | 3 |
| GSN | 1 | Gelsolin | 7 |
| *TUBA1B | 1 | Tubulin alpha chain | 13 |

<sup>#</sup> The protein biomarkers were determined according to the information in the Human Protein Atlas ( https://www.proteinatlas.org/) except three proteins labelled by \*. Those three protein biomarkers were determined based on literature data.

## Discussion

The composition of protein corona directs their biological fates and pharmacokinetics.^11–16^ Proteoforms from the same gene have different biological functions^27–30^ and proteoforms play central roles in modulating disease progression^31–35^. Therefore, characterization of protein corona in a proteoform-specific manner is fundamentally important for improving the overall efficacy of nanomedicine and for discovering novel proteoform biomarkers of diseases from human plasma samples. In this work, the first example of MS-based TDP analysis of NP protein corona was presented to produce fingerprints of proteoforms on the surface of NPs. The TDP technique based on CZE-MS/MS developed in this study offered high-efficient and reproducible proteoform characterization from human plasma PSNP protein corona samples with the detection of hundreds of proteoforms in a mass range of 3-70 kDa, Figures 2 and **4**. The reason of employing CZE-MS/MS instead of commonly used LC-MS/MS for TDP of protein corona is that CZE-MS/MS offers highly efficient separation and highly sensitive detection of proteoforms even large ones.^46,51,52^ The detection of large proteoforms (≥30 kDa) is challenging for typical TDP analysis of complex proteomes due to their substantially lower sensitivity compared to small proteoforms.^50^ Interestingly, clean large proteoform signals were consistently detected for several large proteins (i.e., ∼28, 51, and 66 kDa) from the PSNP protein corona samples, Figure 3. The data demonstrates that coupling CZE-MS/MS-based TDP and protein corona could be a useful strategy for measuring large proteoforms in a complex proteome sample.

Comparing the TDP data with the BUP data of the protein corona sample highlights the value of TDP measurement of intact proteoforms, Figure 4C and **Table 1**. TDP enabled the identification of specific proteoforms with signal peptide cleavage, truncations, and/or PTMs; BUP only provided ambiguous identifications of proteins, typically in the form of protein groups. The MS-based TDP technique developed here will offer much more accurate characterization of protein corona compared to the commonly used BUP approach, which will undoubtedly advance nanomedicine.

The TDP data covers proteoforms of 23 protein biomarkers, *e.g.,* Apolipoproteins, Transthyretin, Complement C3, and Serum amyloid A1. The protein biomarkers in human plasma could exist in multiple different forms, creating a set of proteoforms with potentially heterogenous biological functions. The capability of characterizing protein biomarkers in a proteoform specific manner using TDP will improve the performance of protein biomarkers for disease diagnosis and facilitate the discovery of specific proteoform biomarkers of diseases.

Some additional improvement of the TDP technique will facilitate its broad adoption for NP protein corona analysis. First, more extensive gas-phase fragmentation of proteoforms, especially large ones, will be crucial for the localization of PTMs. Electron- or photon-based fragmentation methods (e.g., electron transfer dissociation,^53^ and ultraviolet photodissociation^54^) need to be integrated into the TDP technique for this purpose. The integration of internal fragment ions^55^ of proteoforms will also be helpful. Second, the numbers of proteoform and proteoform family identifications from the protein corona sample needs to be boosted to cover more low abundance proteoforms to advance nanomedicine and facilitate the discovery of proteoform biomarkers of diseases. Liquid-phase and gas-phase fractionation techniques (e.g., LC^29^ and high field asymmetric waveform ion mobility spectrometry (FAIMS)^56^) can be coupled with CZE-MS/MS to boost the peak capacity of proteoform separations and improve measurement sensitivity, enabling more comprehensive TDP characterization of protein corona.

In summary, this study addresses the limitations of conventional BUP in identifying the precise proteoforms on the protein corona of NPs, which is crucial for predicting their behavior and effectiveness in nanomedicine. The research introduces a novel TDP technique for protein corona analysis, which successfully quantifies hundreds of distinct proteoforms, providing a detailed analysis of proteoforms of over 20 protein biomarkers. The numbers of quantified proteoforms and proteoform families could be significantly increased, using the concept of protein corona sensor array^57^, where multiple NPs are used to increase the depth of proteome. This more accurate characterization of the protein corona using TDP could significantly enhance our understanding of NP biological identities and improve the application of NPs in medicine.

## Methods

### Chemicals and materials

Ammonium bicarbonate (ABC), 3-(trimethoxysilyl) propyl methacrylate, dithiothreitol (DTT), iodoacetamide (IAA), and Amicon Ultra (0.5 mL, 10 kDa cut-off size) centrifugal filter units were ordered from Sigma-Aldrich (St. Louis, MO). LC/ MS grade water, acetonitrile (ACN), HPLC-grade acetic acid (AA), fused silica capillaries (50 mm i.d., 360 mm o.d., Polymicro Technologies) were purchased from Fisher Scientific (Pittsburgh, PA). Acrylamide was obtained from Acros Organics (Fair Lawn, NJ). Healthy human plasma protein was purchased from Innovative Research (www.innov-research.com) and diluted to 55% using phosphate buffer solution (PBS, 1X). Plain Poly Styrene Nanoparticles (PSNPs, ∼100 nm) were obtained from Polysciences (www.polysciences.com).

### Formation of protein corona on the surface of NPs

The protein corona was formed on the PSNPs according to the procedure in a recently published paper^24^ with minor modifications. 75 µL PSNPs (25 mg/mL) were incubated with 55% human plasma (1 mL) for 1 h at 37 °C with constant stirring to form protein coronas. Then, the mixture was centrifugated at 14,000xg for 20 min to remove unbound and loosely attached plasma proteins to the surface of NPs. The protein-NP complexes underwent two cold PBS washes under identical conditions. Subsequently, two-thirds of the resulting protein-NP complexes were collected for the TDP experiment, while the remaining protein-NP complexes were used for the BUP experiment.

### Sample preparation for TDP

The protein-NP complexes were incubated in a 0.4% (w/v) SDS solution for 1.5 hours at 60°C with constant agitation, aiming to elute the protein corona from the surface of the NPs. Subsequently, the supernatant, containing the protein corona in a 0.4% SDS solution, was separated from the PSNPs through centrifugation at 19,000xg for 20 minutes at 4°C. The resulting supernatant underwent an additional centrifugation step under the same condition to ensure complete removal of the PSNPs. The final protein corona sample was cleaned up through a buffer exchange step. An Amicon Ultra Centrifugal Filter with a Molecular Weight Cut-Off (MWCO) of 10 kDa was employed for the buffer exchange, effectively eliminating SDS from the protein samples.

The buffer exchange protocol started with the initial wetting of the filter using 20 µL of 100 mM ammonium bicarbonate (pH 8.0), followed by centrifugation at 14,000xg for 10 minutes. Subsequently, 200 µg of proteins were added to the filter, and centrifugation was carried out for 20 minutes at 14,000xg. 200 µL of 8 M urea in 100 mM ammonium bicarbonate solution was added, followed by centrifugation at 14,000xg for 20 minutes. This step was repeated twice under the same conditions to ensure the complete removal of SDS and other small interferences. To eliminate urea from the purified protein, the filter underwent three additional rounds of buffer exchange. Precisely, 100 mM ammonium bicarbonate was added to the filter, bringing the final volume to 200 µL. All steps were executed with centrifugation at 4°C, ensuring the thorough removal of urea from the protein corona.

After buffer exchange, the concentration of total proteins was determined using a bicinchoninic acid (BCA) kit (Fisher Scientific) in accordance with the manufacturer’s instructions, and the sample was stored at 4°C overnight. The final protein solution, comprising 60 µL of 100 mM ammonium bicarbonate with a protein concentration of 2.4 mg/mL, was collected for CZE-MS/MS analysis.

### Sample preparation for BUP

One-third of the protein corona coated PSNPs prepared in the “Formation of Protein corona on the surface of NPs” section (∼70 μg of total proteins) was dispersed in 35 µL of 100 mM ABC buffer (pH 8.0) containing 8 M urea and led the protein corona to be denatured at 37 °C for 30 min. Then, the protein corona was reduced by adding 5 μL of 70 mM DTT at 37 °C for 30 minutes and alkylated by adding 12.5 μL of 70 mM IAA for 20 minutes in dark at room temperature. The reaction was quenched by adding 1 μL of 70 mM DTT. The protein samples were diluted by four times using 100 mM ammonia bicarbonate, followed by trypsin (1.5 µg, Bovine pancreas TPCK-treated) digestion at 37 °C overnight. The digestion was finally terminated by adding formic acid (0.6 % (v/v) final concentration). The samples were desalted with Sep-Pak C18 Cartridge (Waters, Milford, MA) according to manufacturer’s protocol. The eluates were lyophilized in a vacuum concentrator, and then re-dissolved in 70 µL of 100 mM ABC buffer (pH 8.0).

### CZE-MS/MS

Linear polyacrylamide (LPA)-coated fused silica capillaries (50 µm i.d., 360 µm o.d.) were prepared according to our previous studies.^38,58^ After making the LPA coating, one end of the separation capillary was etched by hydrofluoric acid to reduce its outer diameter to around 100 µm.^59^

The CZE-MS/MS system configuration involved the integration of a CESI 8000 Plus CE system (Beckman Coulter) with an Orbitrap Exploris 480 mass spectrometer (Thermo Fisher Scientific), employing a in-house-built electrokinetically pumped sheath-flow CE-MS nanospray interface.^59,60^ The interface featured a glass spray emitter with an orifice size of 30−35 μm, filled with sheath buffer composed of 0.2% (v/v) formic acid and 10% (v/v) methanol. The spray voltage was about 2 kV. The length of LPA-coated CZE capillary was 100 cm. The capillary’s inlet was securely affixed within the cartridge of the CE system, while its outlet was inserted into the emitter of the interface. The capillary outlet to emitter orifice distance was maintained at approximately 0.5 mm.

For TDP, a 5-psi pressure was applied to load ∼240 ng of corona proteins (2.4 mg/mL, injection volume of 100 nL) into capillary and then, the inlet of the capillary was inserted into the background electrolyte (BGE, 5% (v/v) acetic acid) for CZE separation with a separation voltage of 30 kV. For BUP, 115 nL of each corona peptide sample was loaded for CZE-MS/MS. Following this, the capillary’s inlet was immersed into the BGE (5% (v/v) acetic acid), initiating the CZE separation process under a separation voltage of 30 kV.

For the mass spectrometer, all experiments were conducted using an Orbitrap Exploris 480 mass spectrometer (Thermo Fisher Scientific) in Data-dependent acquisition (DDA) mode. For BUP, full MS scans were acquired in the Orbitrap mass analyzer over the m/z 300-1500 range with resolution of 60,000 (at 200 m/z). Only precursor ions with an intensity exceeding 5E4 and a charge state charge state between 2 and 7 were fragmented in the higher-energy collisional dissociation (HCD) cell and analyzed by the Orbitrap mass analyzer with a resolution of 60,000 (at 200 m/z). One microscan was used. Normalized collision energy was set at 28%. For MS and MS/MS spectra acquisition, the maximum ion injection time was set as 50 and 100 ms, respectively. The precursor isolation width was 2 *m/z*. The dynamic exclusion was applied with a duration of 15 s, and the exclusion of isotopes was enabled.

Two conditions (high-high and low-high) were implemented for the full MS parameters of TDP to enhance precursor ion abundance. In the “high-high” condition, the Full MS parameters included a high mass resolution of 480,000 (at m/z 200) with a single microscan, covering a scan range of 600-2000 m/z. Conversely, the “low-high” condition employed the least Full MS resolution (7,500 at m/z 200) with ten microscans. Other MS parameters remained consistent between the two conditions, encompassing a normalized AGC target value of 300% and an auto maximum injection time. For both conditions, precursor ions in Full MS spectra were isolated with a 2 m/z window and subjected to fragmentation through higher-energy collisional dissociation (HCD) with a normalized collision energy (NCE) of 25%. Only precursor ions with an intensity exceeding 1E4 and a charge state ranging from 5 to 60 were selected for fragmentation. Product ions were detected with a resolution of 120,000 (at 200 m/z), utilizing 3 microscans, and maintaining a normalized AGC target value of 100% for both conditions. Dynamic exclusion was enabled with a duration of 30 seconds and a mass tolerance of 10 ppm (parts per million). Additionally, the “Exclude isotopes” function was activated.

### NP characterization

Dynamic light scattering (DLS) and zeta potential analyses were performed to measure the size distribution and surface charge of the NPs before and after protein corona formation using a Zetasizer nano series DLS instrument (Malvern company). A Helium Neon laser with a wavelength of 632 nm was used for the size distribution measurement at room temperature. Protein corona profiles at the surface of the NPs were studied by sodium dodecyl sulphate–polyacrylamide gel electrophoresis (SDS-PAGE) analysis. Transmission electron microscopy (TEM) was carried out using a JEM-2200FS (JEOL Ltd.) operated at 200kV. The instrument was equipped with an in-column energy filter and an Oxford X-ray energy dispersive spectroscopy (EDS) system. 20 μl of the bare PSNPs deposited onto a copper grid and used for imaging. For protein corona coated NPs, 20 μl of the sample was negatively stained using 20 μl uranyl acetate 1% and finally washed with DI water, deposited onto a copper grid, and used for imaging on the same day.

## Data analysis

For BUP, database searching of the raw files was performed in Proteome Discoverer 2.2 with SEQUEST HT search engine against the UniProt proteome database of human (UP000005640, 82697 entries, version 12/29/2023). Database searching of the reversed database was also performed to evaluate the false discovery rate (FDR). The database searching parameters included full tryptic digestion and allowed up to two missed cleavages, precursor mass tolerance 50 ppm, and fragment mass tolerance 0.05 Da. Carbamidomethylation (C) was set as a fixed modification. Oxidation (M), deamidated (NQ), and acetyl (protein N-term) were set as variable modifications. The data was filtered with a 1% peptide-level FDR. The protein grouping was enabled.

For TDP, proteoform identification and quantification were performed using the TopPIC (Top-down mass spectrometry-based Proteoform Identification and Characterization) pipeline.^43^ In the first step, RAW files were converted into mzML files using the Msconvert tool. The spectral deconvolution which converted precursor and fragment isotope clusters into the monoisotopic masses and proteoform features were then performed using TopFD (Top-down mass spectrometry Feature Detection, version 1.6.3)^61^. The resulting mass spectra and proteoform feature information were stored in msalign and text files, respectively. The database search was performed using TopPIC (version 1.6.3) against a home-built protein database (∼1,000 protein sequences), in which the proteins identified in the BUP data were included. The maximum number of unexpected mass shifts was one. The mass error tolerances for precursors and fragments were 50 ppm. There was a maximum mass shift of 500 Da for unknown mass shifts. To estimate FDRs of proteoform identifications, the target-decoy approach was used and proteoform identifications were filtered by a 1% and 5% FDR at the PrSM level and proteoform level, respectively. The lists of identified proteoforms (TDP) and protein groups (BUP) from all CZE-MS/MS runs are shown in **Supporting Information II**. The TopDiff (Top-down mass spectrometry-based identification of Differentially expressed proteoforms, version 1.6.3) software was used to perform label-free quantification of identified proteoforms by CZE-MS/MS using default settings.^46^

For TDP, the complex sample data was analyzed using Xcalibur software (Thermo Fisher Scientific) to get intensity and migration time of proteoforms. For the final figures, the electropherograms were exported from Xcalibur and formatted using Adobe Illustrator.

## Data Availability

The MS RAW files about TDP measurement were deposited to the ProteomeXchange Consortium via the PRIDE ^62^ with the dataset identifier of PXD050779.

## Supporting information

Proteoform and protein identifications

## Acknowledgments

The authors thank the supports from the National Cancer Institute (NCI) through the grant R01CA247863 (Sun), the National Institute of Diabetes and Digestive and Kidney Diseases (NIDDK) through the grant DK131417 (Mahmoudi), the National Institute of General Medical Sciences (NIGMS) through grants R01GM125991 (Sun) and R01GM118470 (Sun), and the National Science Foundation through the grant DBI1846913 (CAREER Award, Sun).

## Author contributions

S.A.S. performed all the mass spectrometry-related experiments, data analysis, and wrote the draft of the manuscript; A.A.A performed the PSNP characterization including TEM, size, and zeta potential measurements, and helped S.A.S with suggestions regarding the formation of protein corona on PSNP surface. M.M. and L.S. oversaw the project and edited the manuscript.

