## supplementary figures for "Mass spectrometry-based top-down proteomics in nanomedicine: proteoform-specific measurement of protein corona"

* Corresponding Authors.

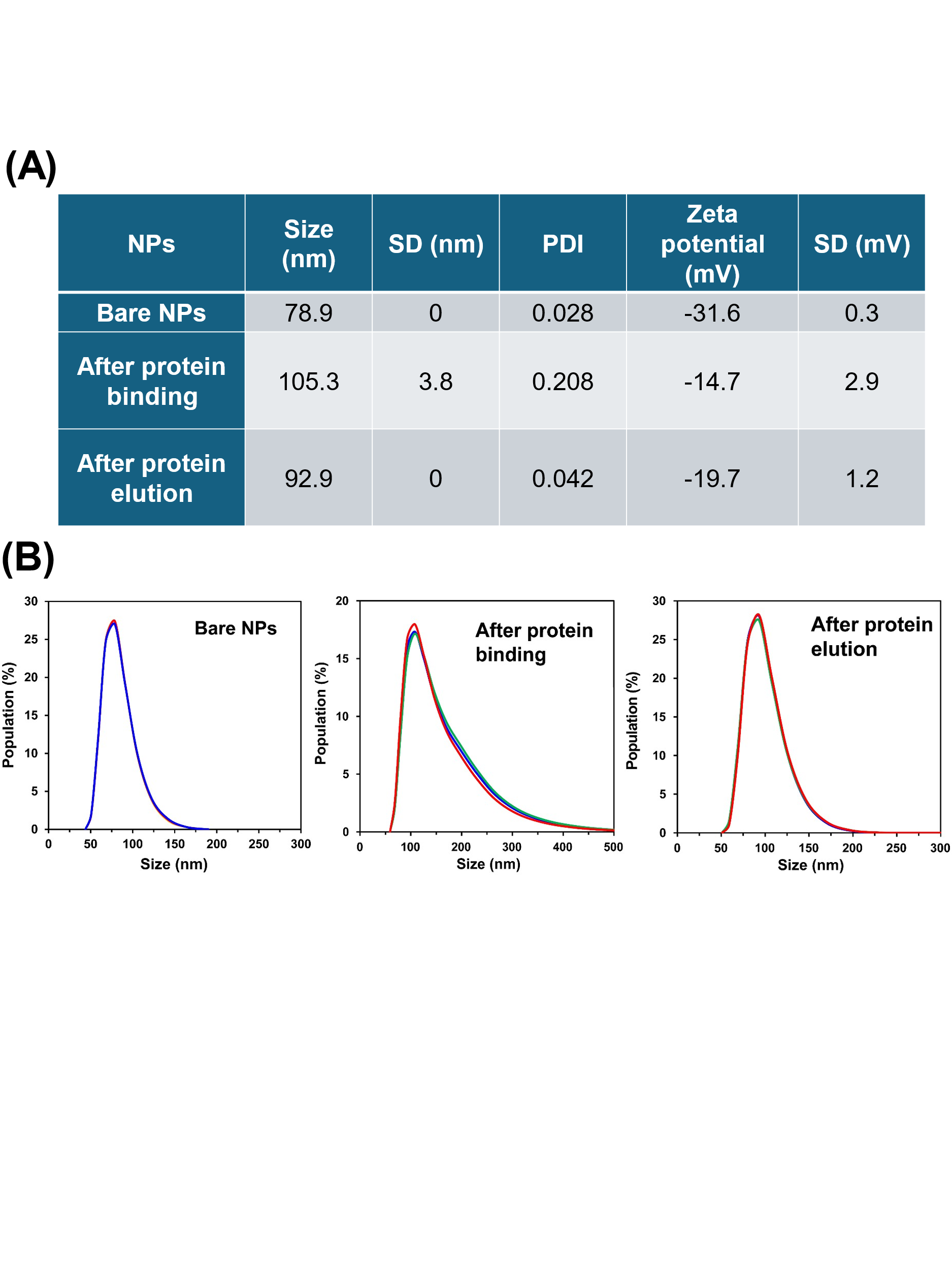

**Figure S1. Characterization of different NP**. (A) Average size, polydispersity index (PDI), zeta potential, and the corresponding standard deviation values of protein corona coated NPs (all measurements are repeated three times, and the corresponding averages are reported). (B) Size distribution and the corresponding replicates of bare and protein corona coated NPs after protein binding and elution, respectively. The results are representative of 3 independent analyses.

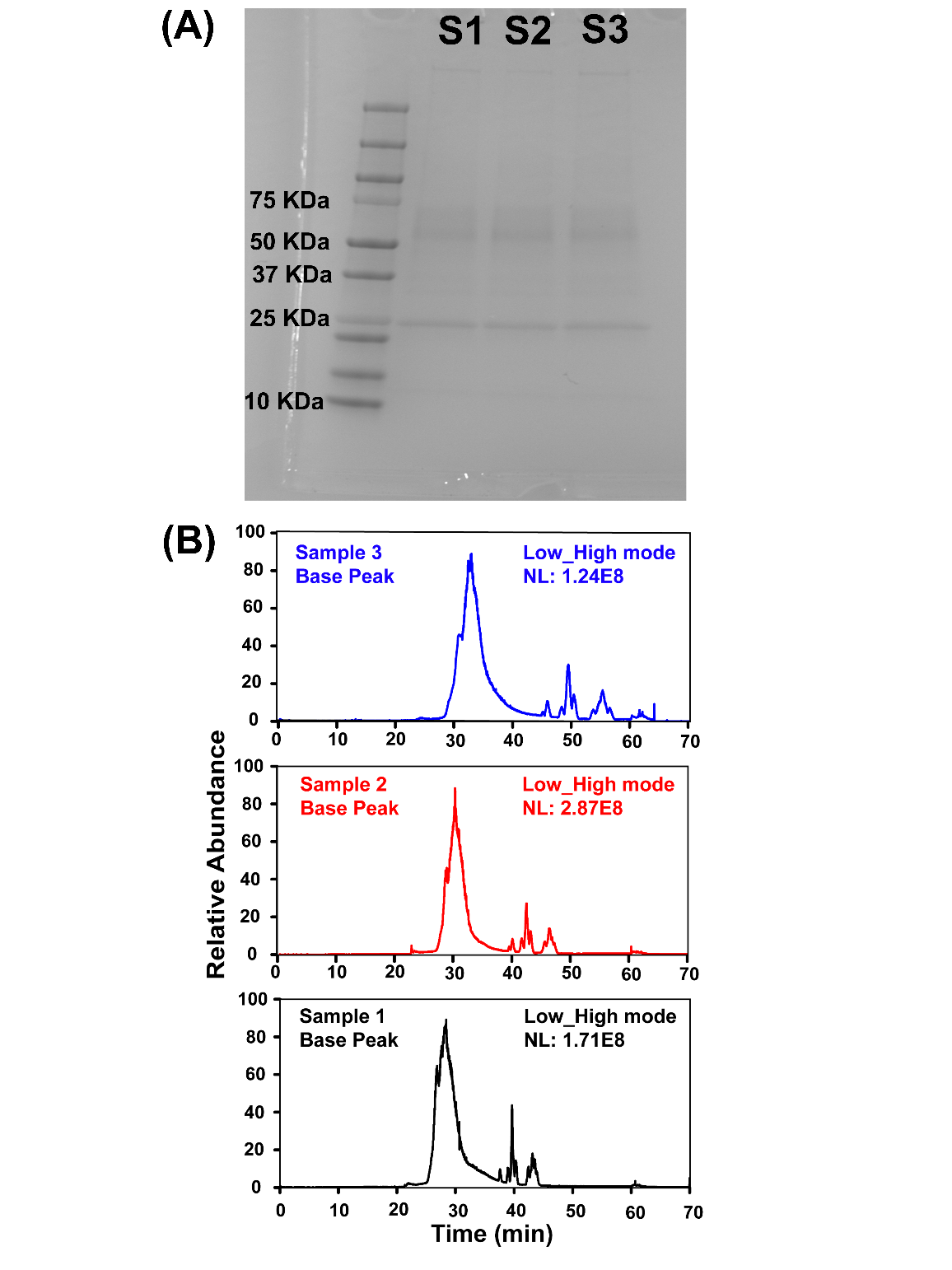

**Figure S2. Reproducibility of CZE-MS/MS for measurements of large proteoforms.** (A) SDS PAGE analysis of three protein corona samples (S1, S2, and S3) prepared in parallel. (B) Base peak electropherograms of the three protein corona samples after CZE-MS/MS analysis in “low-high” mode.

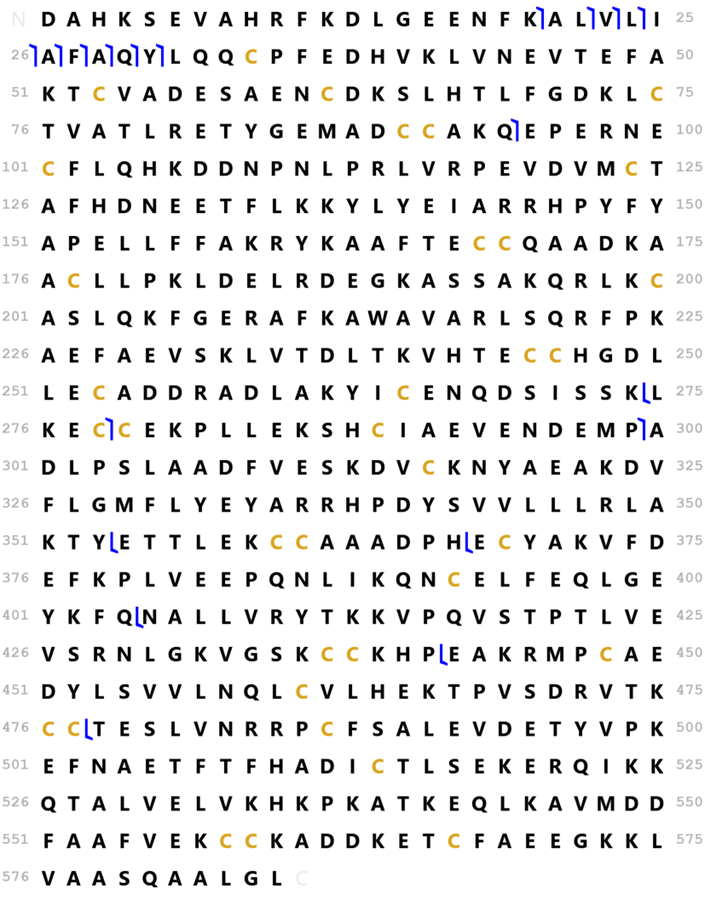

**Figure S3.** Sequence and fragmentation pattern of human serum albumin (HSA) from the CZE-MS/MS analysis (“low-high” mode) of the protein corona sample. The sequence of the mature form of HSA is shown without the signal peptide and the propeptide. No post-translational modifications (PTMs) and disulfide bonds were considered. The ProSight Lite software was used to match the experimental MS/MS data with the target protein sequence with a 50-ppm mass tolerance. The mass tolerance was determined based on the mass error of our instrument when the experiment was done.

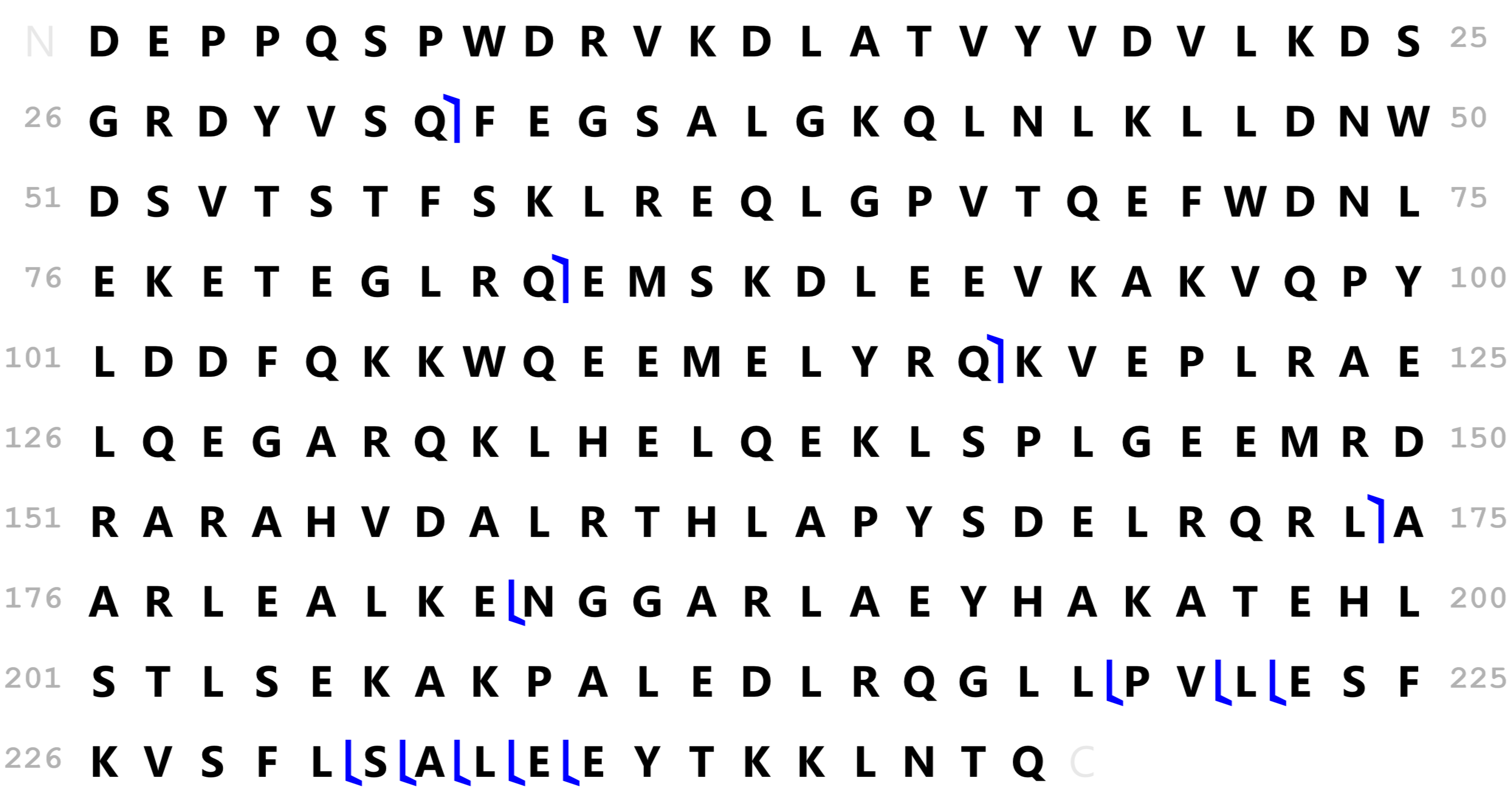

**Figure S4**. Sequence and fragmentation pattern of Apolipoprotein A-I (APOA1) from the CZE-MS/MS analysis (“low-high” mode) of the protein corona sample. The sequence of the mature form of APOA1 is shown without the signal peptide. No post-translational modifications (PTMs) were considered. The ProSight Lite software was used to match the experimental MS/MS data with the target protein sequence with a 50-ppm mass tolerance. The mass tolerance was determined based on the mass error of our instrument when the experiment was done.
